# The U2AF homology motif kinase 1 (UHMK1) is upregulated upon hematopoietic cell differentiation

**DOI:** 10.1101/187385

**Authors:** Isabella Barbutti, João Agostinho Machado-Neto, Vanessa Cristina Arfelli, Paula de Melo Campos, Fabiola Traina, Sara Teresinha Olalla Saad, Leticia Fröehlich Archangelo

**Author notes:** Corresponding author: Department of Cellular and Molecular Biology and Pathogenic, Bioagents, Ribeirão Preto Medical School, University of São Paulo, Av Bandeirantes 3900, 14049-900 Ribeirão Preto-SP, Brazil.; (L.F. Archangelo).

## Abstract

UHMK1 (KIS) is a nuclear serine/threonine kinase that possesses a U2AF homology motif and phosphorylates and regulates the activity of the splicing factors SF1 and SF3b155. Mutations in these components of the spliceosome machinery have been recently implicated in leukemogenesis. The fact that UHMK1 regulates these factors suggests that UHMK1 might be involved in RNA processing and perhaps leukemogenesis. Here we analyzed UHMK1 expression in normal hematopoietic and leukemic cells as well as its function in leukemia cell line.

In the normal hematopoietic compartment, markedly higher levels of transcripts were observed in differentiated lymphocytes (CD4^+^, CD8^+^ and CD19^+^) compared to the progenitor enriched subpopulation (CD34^+^) or leukemia cell lines. *UHMK1* expression was upregulated in megakaryocytic-, monocytic-and granulocytic-induced differentiation of established leukemia cell lines and in erythrocytic-induced differentiation of CD34^+^ cells. No aberrant expression was observed in patient samples of myelodysplastic syndrome (MDS), acute myeloid (AML) or lymphoblastic (ALL) leukemia. Nonetheless, in MDS patients, increased levels of *UHMK1* expression positively impacted event free and overall survival.

Lentivirus mediated UHMK1 knockdown did not affect proliferation, cell cycle progression, apoptosis or migration of U937 leukemia cells, although UHMK1 silencing strikingly increased clonogenicity of these cells. Thus, our results suggest that UHMK1 plays a role in hematopoietic cell differentiation and suppression of autonomous clonal growth of leukemia cells.

## 1. Introduction

UHMK1 was first identified as a kinase that interacts with Stathmin (KIS) [1], an important regulator of microtubule dynamics. Since then it has been described to interact with a range of proteins, such as peptidylglycine α-amidating monooxygenase (PAM) [2]; cyclin dependent kinase inhibitor (CDKI) p27^KIP1^ [3]; splicing factors SF1 [4] and SF3b155 [5]; components of the neuronal RNA granules NonO, KIF3A and eEF1A [6]; proliferation marker PIMREG [7]; and RNA-binding proteins CPEB1, CPEB2 and CPEB3 [8], shedding light on different functions of UHMK1 in diverse cellular processes.

UHMK1 is ubiquitously, but preferentially, expressed in neural systems [9], its mRNA levels increase gradually during postnatal development, reaching highest levels in the mature brain. Not surprisingly, most of UHMK1-related functions have been addressed in the context of adult nervous system [6, 8, 10-18].

Other than the neuronal milieu, important functions of UHMK1 are conferred to its unique feature. UHMK1 is the only known kinase to possess the N-terminal kinase core juxtaposed to a C-terminal U2AF homology motif (UHM), which shares significant similarity to the SF1-binding UHM domain of the splicing factor U2AF^65^ [19].

Through the UHM motif, UHMK1 is capable of interacting with splicing factors such as SF1 and SF3b155 [5]. Upon interaction, UHMK1 phosphorylates SF1, which in turn favors the formation of a U2AF^65^-SF1-RNA complex at 3′ splice site, an event known to take place during the early steps of spliceosome assembly [4]. Moreover, UHMK1 expression is necessary for normal phosphorylation of SF1 *in vivo* [16]. Thus UHMK1 may participate in RNA splicing.

Another important function described for UHMK1 is its ability to positively regulate cell cycle progression. Upon mitogen activation, UHMK1 is upregulated and phosphorylates p27^Kip1^ on serine 10 (S10), promoting its nuclear export and release of its inhibitory effect on cell cycle [3]. Therefore, UHMK1 promotes cell cycle re-entry by inactivating p27^Kip1^ following growth factor stimulation, as demonstrated during proliferative response of vascular smooth cells (VSMC) [20] and corneal endothelial cells (CEC) [21, 22]. Accordingly, deletion of UHMK1 abolished VSMC proliferation whereas it increased its migratory activity [23].

Thus abnormally elevated UHMK1 activity, which is expected to relieve cells from growth inhibition dependent on p27^Kip^, could be involved in some aspects of tumor development. Indeed, Zhang and coworkers demonstrated that the UHMK1 silencing enhances anti-tumor activity of erlotinib and could be a molecular target for combinatorial treatment in EGFR TKI-resistant cases of breast cancer [24]. More recently, UHMK1 was described in a panel of novel serum tumor antigen-associated autoantibody (TAAb) biomarkers for detection of serous ovarian cancer [25].

We described UHMK1 interaction with PIMREG (previously known as CATS; FAM64A), a protein highly expressed in leukemia cells, know to interact with and influence the subcellular localization of the leukemic fusion protein PICALM/MTT10 [26]. We suggested a possible role of UHMK1-PIMREG interaction in PICALM/MTT10 mediated leukemogenesis [7].

Interestingly, an increasing amount of data has pointed to altered splicing machinery as a novel mechanism in leukemogenesis [27]. Somatic mutation has been found in genes coding for splicing factors in patients with diverse hematological malignancies, including the UHMK1 substrates SF1 and SF3b155 [28, 29].

The described role for UHMK1 in controlling cell cycle progression, its interaction with a proliferation marker and most importantly is function in regulation of the splicing factors commonly found mutated in hematological malignancies, prompted us to investigate a possible function of UHMK1 in the hematopoietic compartment and possibly leukemogenesis.

Here we show that UHMK1 expression is upregulated upon differentiation of hematopoietic cells. Of note, depletion of its protein does not interfere with proliferation or cell cycle progression of leukemic cells but rather it increases the clonogenicity of the U937 leukemia cell line.

## 2. Materials and methods

### 2.1. Sorting of human hematopoietic cell subpopulation

For sorting of human cells, peripheral blood samples obtained from healthy donors were treated with lysis buffer containing ammonium chloride (red blood cells lysis). Samples underwent density-gradient separation through Ficoll-Paque Plus (GE Healthcare, Uppsala, Sweden) according to the manufacturer’s instructions.

Mononuclear cells were isolated and subsequently incubated for 30 min on ice with phycoerythrin (PE)-labeled CD19 (B lymphocytes), allophycocyanin (APC)-labeled CD8 (T lymphocytes), fluorescein isothiocyanate (FITC)-labeled CD4 (T lymphocytes) and peridinin-chlorophyll protein complex (PerCP)-labelled CD3. Cells were sorted using FACSAria™ IIu (BD Biosciences, Palo Alto, CA). Granulocytes were obtained from the Ficoll-Paque Plus polymorphonuclear cell fraction.

For CD34^+^ cell isolation, mononuclear cells were labeled with CD34 microbeads for 40 min at 4°C and isolated using MACS magnetic cell separation columns (Miltenyi Biotec, Mönchengladbach, Germany) according to the manufacturer’s instructions.

All cells were subsequently used for RNA extraction.

### 2.2. Patient samples

Total bone marrow samples obtained from healthy donors (n=19), MDS (n=45), AML (n=42) and ALL (n=22) patients were analyzed. All patients included in the study were untreated at the time of sample collection. Patients’ characteristics are depicted in Table 1. The study was approved by the local ethics committee of University of Campinas and all subjects provided written informed consent. Patients that attended the clinic between 2005 and 2016 and signed their consent were included. Acute promyelocytic leukemia diagnoses were not included in the study. MDS patients were classified according to WHO 2008 classification [30], and cytogenetic risk for AML was defined according to British Medical Research Council (MRC) category [31].

**Table 1.**
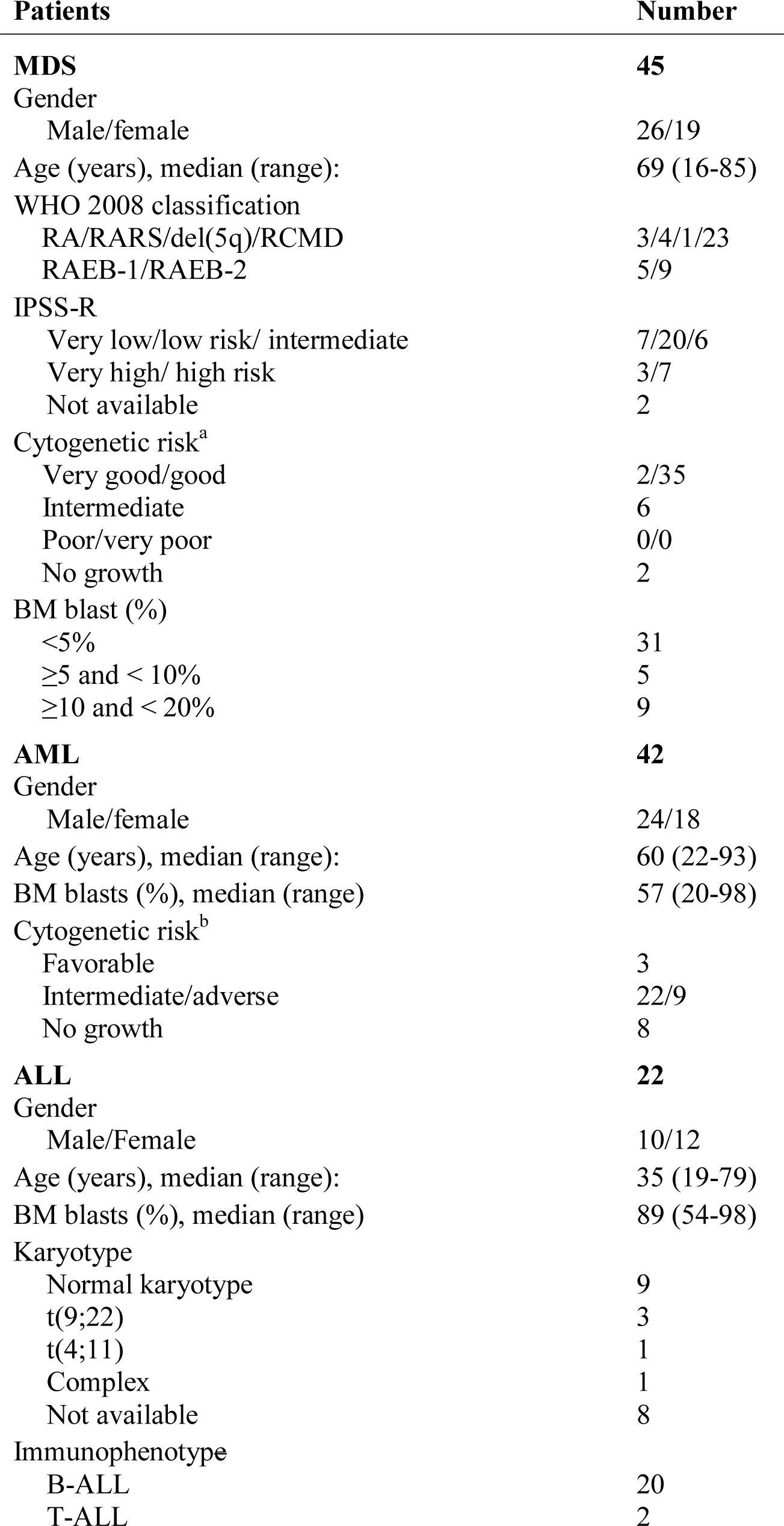
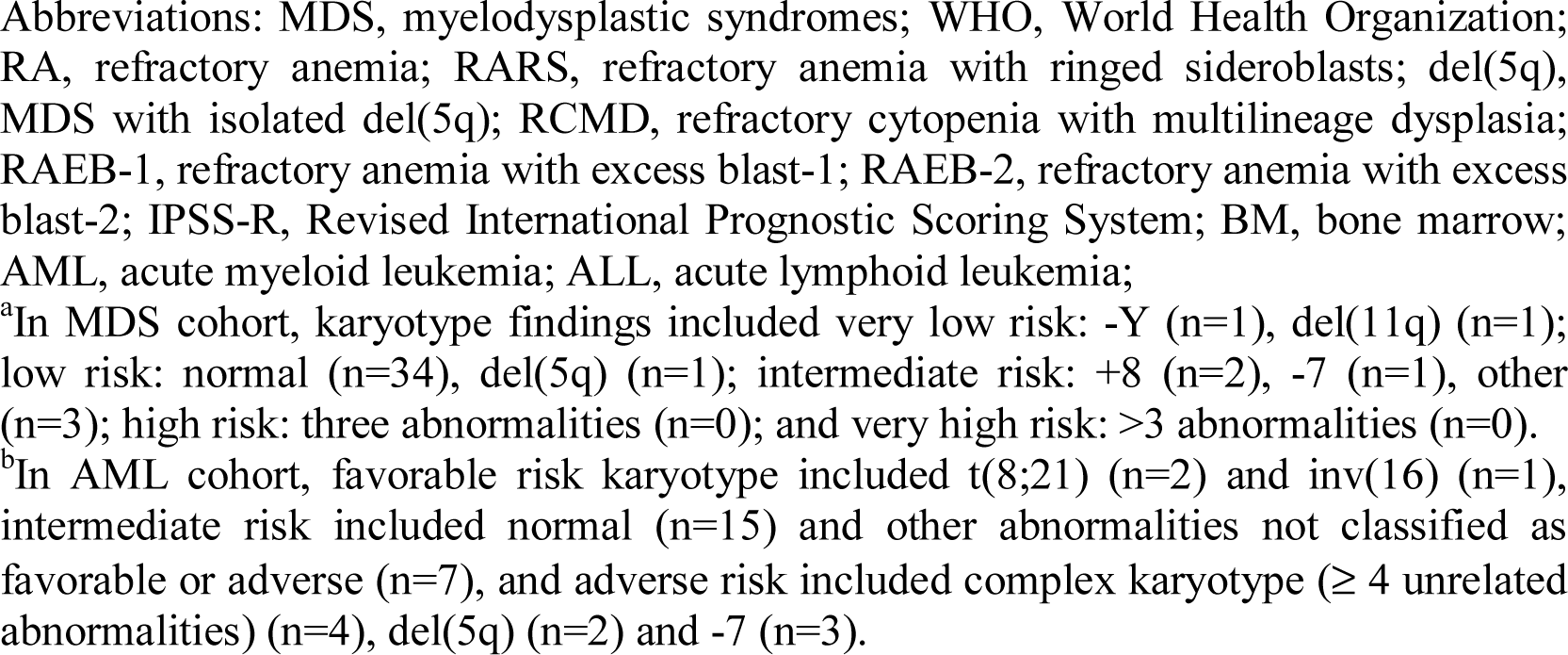
Patients’ characteristics

| **Patients** | **Number** |
| --- | --- |
| **MDS** | **45** |
| Gender |  |
| Male/female | 26/19 |
| Age (years), median (range): | 69 (16-85) |
| WHO 2008 classification |  |
| RA/RARS/del(5q)/RCMD | 3/4/1/23 |
| RAEB-1/RAEB-2 | 5/9 |
| IPSS-R |  |
| Very low/low risk/ intermediate | 7/20/6 |
| Very high/ high risk | 3/7 |
| Not available | 2 |
| Cytogenetic risk<sup>a</sup> |  |
| Very good/good | 2/35 |
| Intermediate | 6 |
| Poor/very poor | 0/0 |
| No growth | 2 |
| BM blast (%) |  |
| <5% | 31 |
| ≥5 and < 10% | 5 |
| ≥10 and < 20% | 9 |
| **AML** | **42** |
| Gender |  |
| Male/female | 24/18 |
| Age (years), median (range): | 60 (22-93) |
| BM blasts (%), median (range) | 57 (20-98) |
| Cytogenetic risk<sup>b</sup> |  |
| Favorable | 3 |
| Intermediate/adverse | 22/9 |
| No growth | 8 |
| **ALL** | **22** |
| Gender |  |
| Male/Female | 10/12 |
| Age (years), median (range): | 35 (19-79) |
| BM blasts (%), median (range) | 89 (54-98) |
| Karyotype |  |
| Normal karyotype | 9 |
| t(9;22) | 3 |
| t(4;11) | 1 |
| Complex | 1 |
| Not available | 8 |
| Immunophenotype |  |
| B-ALL | 20 |
| T-ALL | 2 |

Abbreviations: MDS, myelodysplastic syndromes; WHO, World Health Organization; RA, refractory anemia; RARS, refractory anemia with ringed sideroblasts; del(5q), MDS with isolated del(5q); RCMD, refractory cytopenia with multilineage dysplasia; RAEB-1, refractory anemia with excess blast-1; RAEB-2, refractory anemia with excess blast-2; IPSS-R, Revised International Prognostic Scoring System; BM, bone marrow; AML, acute myeloid leukemia; ALL, acute lymphoid leukemia;
<sup>a</sup>In MDS cohort, karyotype findings included very low risk: -Y (n=1), del(11q) (n=1); low risk: normal (n=34), del(5q) (n=1); intermediate risk: +8 (n=2), -7 (n=1), other (n=3); high risk: three abnormalities (n=0); and very high risk: >3 abnormalities (n=0).
<sup>b</sup>In AML cohort, favorable risk karyotype included t(8;21) (n=2) and inv(16) (n=1), intermediate risk included normal (n=15) and other abnormalities not classified as favorable or adverse (n=7), and adverse risk included complex karyotype ( $\geq 4$ unrelated abnormalities) (n=4), del(5q) (n=2) and -7 (n=3).

### 2.3. Leukemia cell lines

A panel of leukemia cell lines including myeloid (U937, K562, NB4 and KU812) and lymphoid (Jurkat, MOLT-4 and Namalwa) cells were obtained from the American Type Culture Collection (ATCC, Manassas, USA) or the German Collection of Microorganisms and Cell Cultures (DSMZ, Braunschweig, Germany), grown according to the suppliers’ recommendations and used for RNA extraction, cell differentiation and/or virus transduction. All cell lines were tested and authenticated by STR matching analysis using the PowerPlex^®^ 16 HS system (Promega, Madison, WI, USA) and the ABI 3500 Sequence Detector System (Applied Biosystems, Foster City, CA, USA).

### 2.4. Differentiation of cell lines and primary CD34^+^ cells

RNA and protein samples of leukemia cells and primary peripheral blood CD34^+^ cells induced to differentiate were obtained from previous studies [32, 33]. *UHMK1* expression was assessed in Hemin and Hydroxyurea (HE-HU)-induced erythroid differentiation of KU812 (n=3), phorbol-13 myristate-12 acetate (PMA)-induced megakaryocytic differentiation of K562 (n=3), all-trans retinoic acid (ATRA)-induced granulocytic differentiation of NB4 (n=6), ATRA-induced granulocytic differentiation of U937 (n=4) and PMA-induced monocytic differentiation of U937 (n=4). Progenitor and differentiated erythrocytes were obtained from CD34^+^ derived BFU-E, CFU-E and pro-erythroblasts (6 days) cultured for further 6 days with erythropoetin (EPO) and holo-transferin (12 days) (n=6).

### 2.5. Peripheral blood lymphocytes (PBLs) activation

Human resting PBLs were prepared from peripheral blood of healthy donors by standard Ficoll/Hypaque gradient centrifugation and stimulated with phytohemaglutinin (PHA) as previously described [34].

### 2.6. Quantitative PCR

Total RNA was extracted using Trizol^®^ or RNeasy^®^ Mini or Micro Kit (Qiagen, Hilden, Germany). DNAse I treated RNA was reverse transcribed with oligo dT primers and RevertAid™ First Strand cDNA Synthesis Kit (MBI Fermentas, St. Leon-Rot, Germany). Reactions were carried out with Maxima SYBR Green qPCR master mix, according to the manufacture’s protocol (MBI Fermentas). Plates were run and analyzed using the 7500 RealTime PCR Systems (Life Technologies, Carlsbad, CA, USA). *HPRT* and *GAPDH* were used as endogenous controls, and relative gene expression was calculated by 2^−ΔΔCt^ equation [35].

### 2.7. Immunoblotting

Cellular lysates were electrophoresed on 10-12% SDS-PAGE and transferred to nitrocellulose membrane (Hybond^TM^ ECL^TM^, GE Healthcare, Buckinghamshire, UK). The membranes were blocked and probed with specific antibody, followed by detection with fluorescently labeled secondary antibodies. Membranes were visualized on Alliance 2.7 (UVItec, Cambridge, England). Primary antibodies were anti-UHMK1 (KIS3B12; 1:10) [16], anti-KIST mAb (Abcam-117936; 1:1000), anti-p27 (1:1000) (sc-1641, Santa Cruz Biotechnologies), anti-phospho-p27(S10) (1:1000) (34-6300, ThermoFisher), anti-GAPDH (1:4000) (sc-32233, Santa Cruz Biotechnologies) and anti-β-ACTIN (1:20000) (A5441, Sigma). Secondary antibodies were purchased from Life Technologies (Carlsbad, CA, USA): anti-mouse (1:5000) and anti-rat (1:2000).

### 2.8. Transduction of U937 cells

Monocytic U937 cells were transduced with lentiviruses particles expressing a pool of three different short hairpin RNAs (shRNA) targeting the *UHMK1* sequence (UHMK1 sc-78640-V; Santa Cruz Biotechnologies, CA, USA; siRNA target sequences CCAGAAGCAGAAUUGCAAAtt, GGUUCUUCCUUAUUGUUGAtt, and GAUGCUUGAUCUUGCACAAtt) or nonspecific control target (sc-108080), named shUHMK1 and shControl cells, respectively.

Briefly, 2 x 10^5^ cells were transduced by spinoculation at multiplicity of infection (MOI) equal to 1. Stable polyclonal shUHMK1 and shControl cell lines were stablished after 15 days of selection with appropriate antibiotics (1 μg/mL).

### 2.9. Methylthiazoletetrazolium (MTT) assay

For MTT assay cells were seeded in 96-well plates at density of 2.5 x 10^4^ cells/well. After 48 hours of normal culture conditions, 10 μL of a 5 mg/mL solution of MTT was added per well and incubated for 4 hours. The reaction was stopped by adding 100 μL of 0.1 N HCl in anhydrous isopropanol. Cell viability was evaluated by measuring the absorbance at 570 nm with an automated plate reader.

### 2.10. Cell cycle assay

2.5 x 10^5^ cells were fixed in 70% ethanol for at least 30 min at 4°C, washed with PBS and stained with 20 μg/mL propidium iodide (PI) containing 10 μg/mL RNAse A for 30 min at room temperature (RT). Cell cycle analysis was performed using FACSCalibur (Becton-Dickinson, California, USA) and Modfit (Verify Software House Inc., USA).

### 2.11. Apoptosis assays

Cells were washed with ice-cold PBS and stained with annexin V and PI (BD Biosciences Pharmingen, California, USA) for 15 minutes at RT. Apoptosis analysis was performed using FACSCalibur (Becton-Dickinson) and FACSDiva software (BD Biosciences Pharmingen). Ten thousand events were acquired for each sample.

### 2.12. Migration assay

Migration assay was performed in a 5μm pore-sized transwell plates (Costar, Corning, NY, USA). Cells were seeded above the filters at a density of 1 x 10^5^ cells/well. The lower compartment was filled with the following media: 10% FBS/0.5% BSA; 200 ng/mL CXCL12/0.5% BSA (PeproTech, Rocky Hill, NJ, USA) and 0.5% BSA, was used as negative control. After 24 hours, the number of cells which migrated through the filter and reached the lower compartment was counted. Values were expressed as percentage of the input (cells applied directly to the lower compartment) set as 100%.

### 2.13. Colony forming assay

shUHMK1 and shControl transduced U937 cells were plated in semisolid medium depleted of any growth factors (5 x 10^2^ cell/ml; MethoCult 4230; StemCell Technologies Inc., Vancouver, Canada). Colonies were detected after 8 days of culture by adding 200 μl of a 5 mg/ml MTT solution and scored by Image J quantification software (NIH, Bethesda, Maryland, USA).

### 2.14. Statistical analysis

Statistical analysis was performed using GraphPad Instat Prism 5 (GraphPad Software Inc., San Diego, CA, USA) and the Statistical Analysis System (SAS) for Windows 9.4 (SAS Institute Inc, Cary, NC, USA). Two-tailed Student’s t test was used for paired measured factors. Results are shown as mean ± standard deviation of mean (SEM). Cox regression model was used to estimate overall survival (OS) and event-free survival (EFS) for patients with MDS. The stepwise process of selection was used for multivariate analysis. OS was defined from the time of sampling to the date of death or last follow-up, and EFS was defined as the time of sampling to the date of progression to high-risk MDS or AML with myelodysplasia-related changes, date of death or last follow-up. A *p* value <0.05 was considered as statistically significant.

## 3. Results

### 3.1. *UHMK1* expression in hematopoietic subpopulations

QPCR was performed in order to determine *UHMK1* expression in the hematopoietic compartment. Expression analysis revealed high and variable levels of *UHMK1* in the hematopoietic subpopulations and leukemia cell lines.

Markedly higher expression of *UHMK1* was observed in differentiated T-lymphocytes (CD4^+^ and CD8^+^) and B cells (CD19^+^), with an increase of over 50-fold changes compared to bone marrow sample (BM) (Figure 1A). Elevated levels of transcripts were observed in the chronic myeloid leukemia KU812 cell line with 4-fold changes expression compared to BM or higher when compared to other leukemia cell lines, such as U937, K562, NB4, Jurkat, MOLT-4 and Namalwa (Figure 1B).

**Figure 1:**
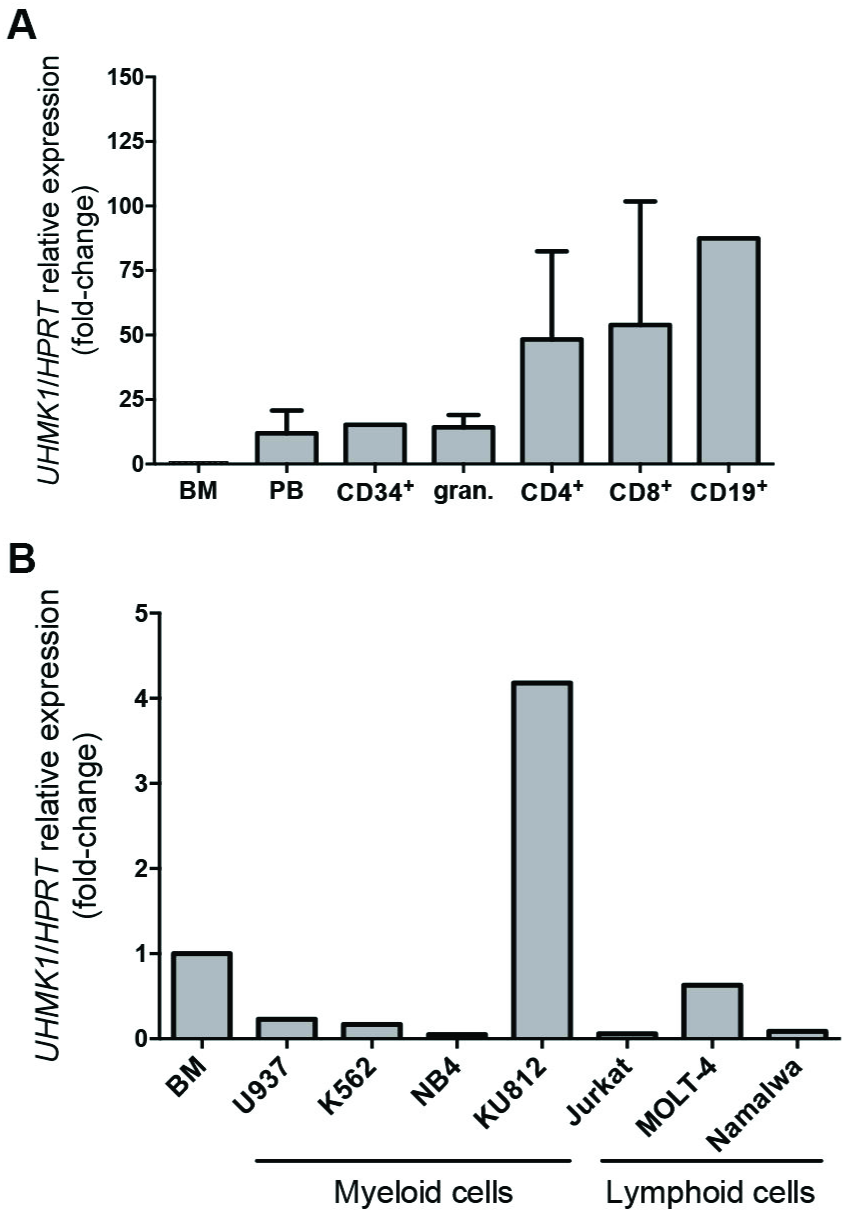
*UHMK1*expression in normal hematopoietic subpopulations and leukemia cell lines. Relative *UHMK1* expression normalized by *HPRT*. Samples are: whole bone marrow (BM), **(A)** peripheral blood (PB), progenitor cells (CD34^+^), granulocytes (gran.), T-lymphocytes (CD4^+^ and CD8^+^) and B-lymphocytes (CD19^+^), **(B)** myeloid leukemia cell lines (U937, K562, NB4 and KU812) and lymphoid leukemia cell lines (Jurkat, MOLT-4 and Namalwa).

### 3.2. UHMK1 expression is upregulated during induced differentiation of hematopoietic cells

UHMK1 gene and protein expression were evaluated during differentiation of established leukemia cell line models. Levels of *UHMK1* mRNA increased by 2-fold changes during megakaryocytic, by 8-fold changes during monocytic and up to 6-fold changes during granulocytic differentiation of both NB4 and U937 cells (Figure 2). The increase in mRNA expression was mostly accompanied by protein accumulation; except in granulocyte differentiated NB4 cells (Figure 2D). UHMK1 expression was not altered in KU812-differentiated erythrocytes, in comparison to the same non-induced cells (Figure 2A). Further, *UHMK1* expression was analyzed in primary peripheral blood CD34^+^ cells induced to differentiate into erythrocytes *in vitro* (n=6)*. UHMK1* was upregulated by 4-fold changes on differentiated erythrocytes (12 days) in comparison to undifferentiated cells (6 days) (Figure 2F).

**Figure 2:**
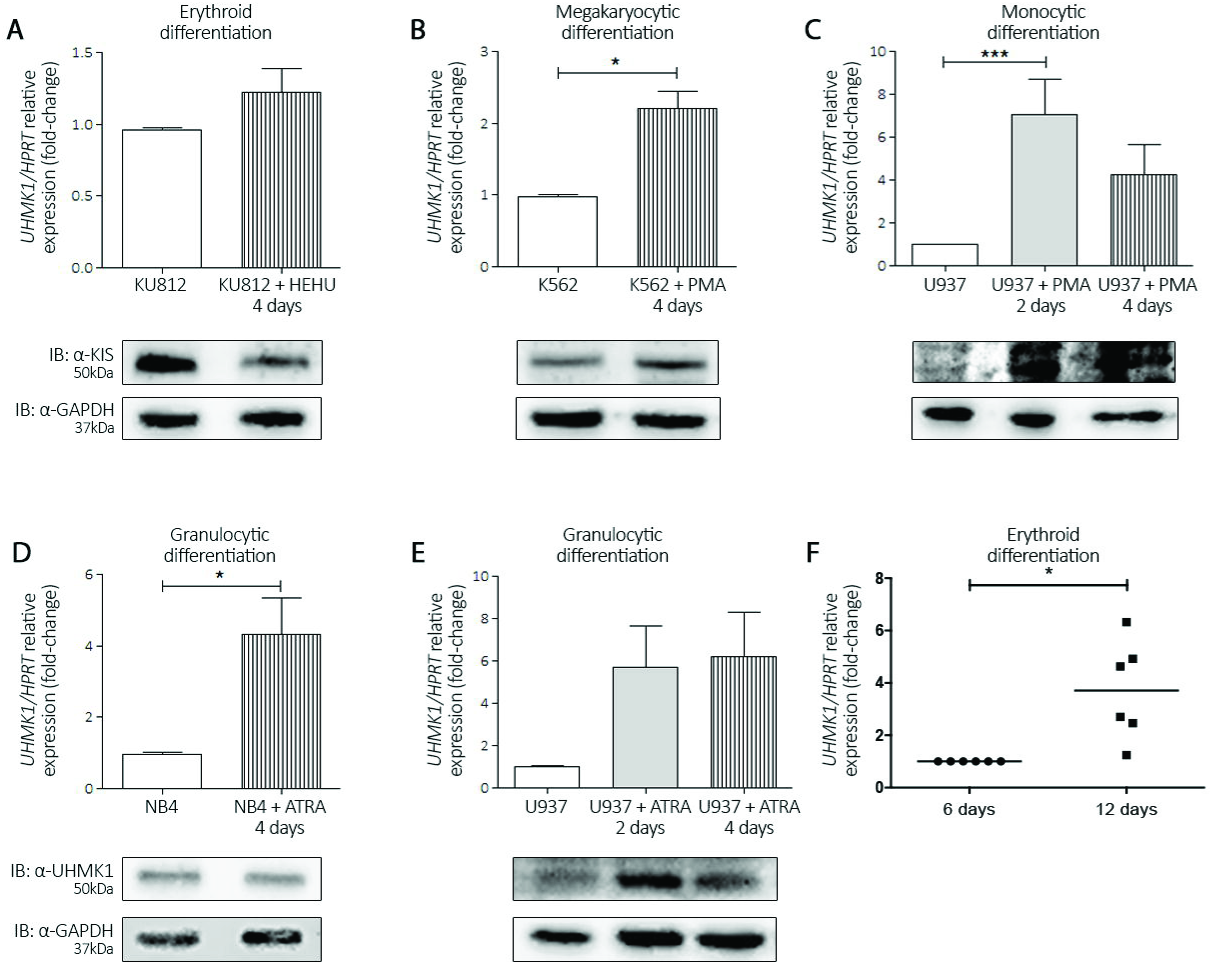
*UHMK1* mRNA and protein expression during induced differentiation of leukemia cell lines and primary peripheral blood CD34^+^ erythroid cell differentiation. Relative *UHMK1* expression at days 2 and/or 4 of differentiation. *UHMK1* mRNA levels normalized by *HPRT* are shown in upper panels. Western blot of total cell extracts blotted against anti-UHMK1 and anti-GAPDH are shown in lower panels. Results are shown as mean ± Standard Deviation (SD) of at least three independent experiments. **(A)** HE+HU-induced erythroid differentiation of KU812 cells. **(B)** PMA-induced megakaryocytic differentiation of K562 cells. **(C)** PMA-induced monocytic differentiation of U937 cells. **(D)** ATRA-induced granulocytic differentiation of NB4 (left panel) and U937 cells (right panel). \**p*<0.05; \*\**p*<0.01 and \*\*\**p*<0.001, Student’s t test. **(F)** Relative *UHMK1* expression in progenitor (6 days) and differentiated (12 days) erythrocytes obtained from erythropoetin (EPO) and holo-transferin cultured primary peripheral blood CD34^+^ cells (n=6). *UHMK1* expression was normalized by *HPRT*. \**p*=0.0169; Student’s t test.

### 3.3. *UHMK1* expression in MDS and acute leukemia patient samples

Analysis of *UHMK1* expression was evaluated by qPCR in BM samples of MDS and acute leukemia patients. No aberrant expression of *UHMK1* transcript was observed comparing healthy donors [median: 1.0 (range: 0.07-3.27)], MDS [0.67 (0.06-2.51)], AML [0.88 (0.02-4.58) and ALL [0.54 (0.05-7.62); all *p*>0.05) patients. Similarly, no differences in *UHMK1* expression were observed when MDS patients were stratified by WHO 2008 classification into RA/RARS/del(5q)/RCMD and RAEB-1/RAEB-2 groups and AML patients according to cytogenetic risk stratification (Figure 3).

**Figure 3:**
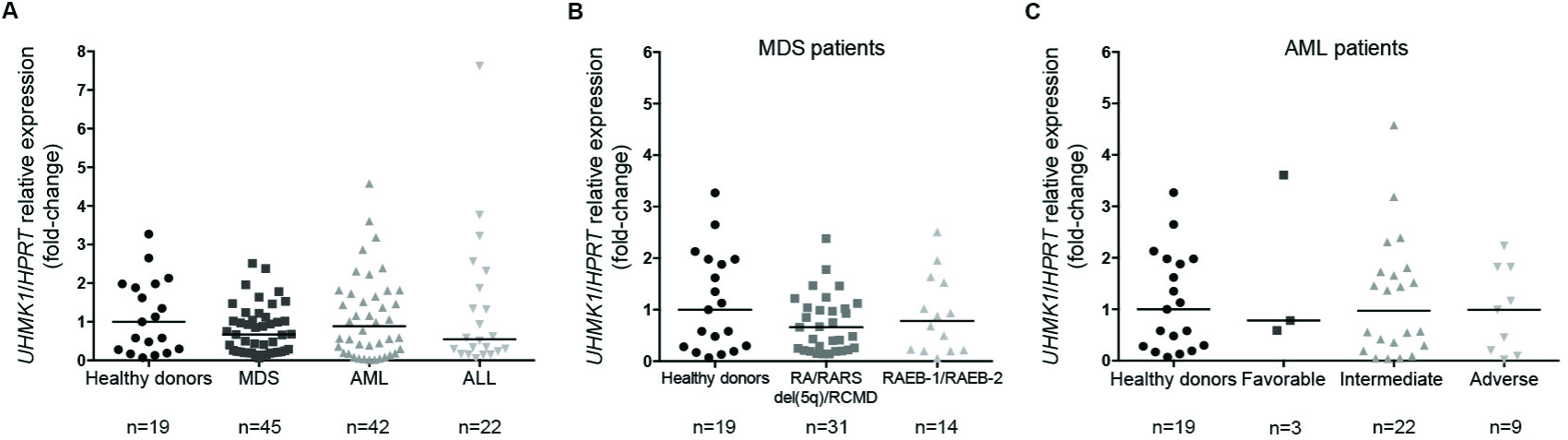
*UHMK1* expression in MDS and leukemia patient samples. qPCR analysis of *UHMK1* mRNA levels in **(A)** total bone marrow cells of healthy donors and patients with MDS, AML and ALL; **(B)** MDS patients stratified by WHO 2008 classification; **(C)** AML patients according to cytogenetic risk stratification. Horizontal lines represent the median; all *p* values were above 0.05 (Mann-Whitney test).

Further, the association of *UHMK1* expression with known clinical factors linked with MDS progression was analyzed. In our cohort of MDS patients, clinical factors that significantly affected both event free and overall survival included WHO 2008 diagnosis (RAEB-1/RAEB-2 vs. others, *p*<0.0001) and IPSS-R (very high/high/intermediate *vs.* very low/low, *p*<0.0002) by univariate analysis. Multivariate analyses indicated that increased levels of *UHMK1* expression positively impacted event free survival (hazard ratio: 0.527; 95% C.I.: 0.281-0.989; *p* = 0.0431) and overall survival (hazard ratio: 0.522; 95% C.I.: 0.275-0.992; *p* = 0.0437) along with RAEB-1/RAEB-2 WHO 2008 classification (*p*<0.0001) (Table 2).

**Table 2.**
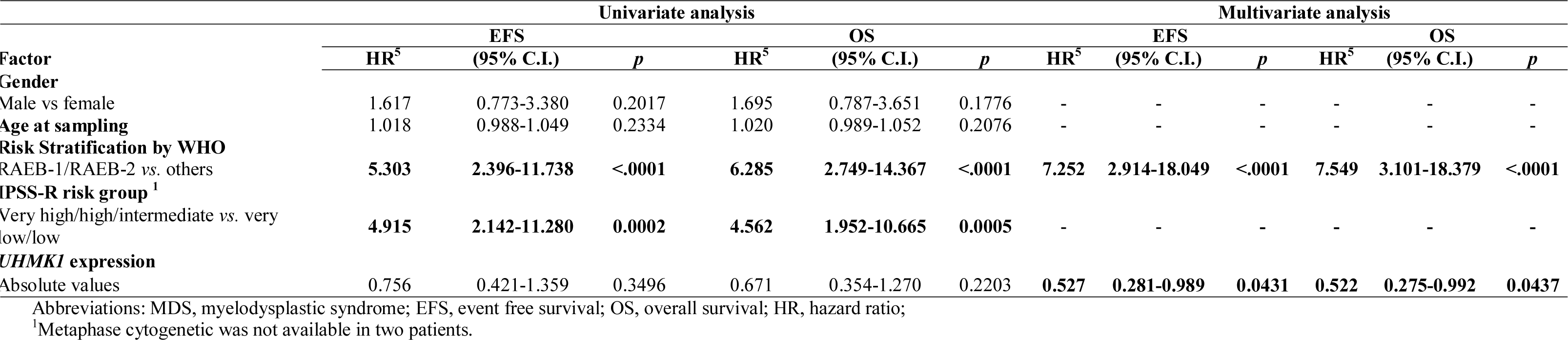
Univariate and multivariate analyses of survival outcomes for MDS patients

### 3.4. UHMK1 silencing does not affect cell cycle progression of U937 leukemia cells

The U937 leukemia cell line was transduced with lentiviral particles expressing a pool of multiple shRNA targeting the *UHMK1* mRNA. UHMK1 was efficiently depleted at both mRNA and protein levels in shUHMK1 transduced cells compared to shControl cells (Figure 4A).

**Figure 4:**
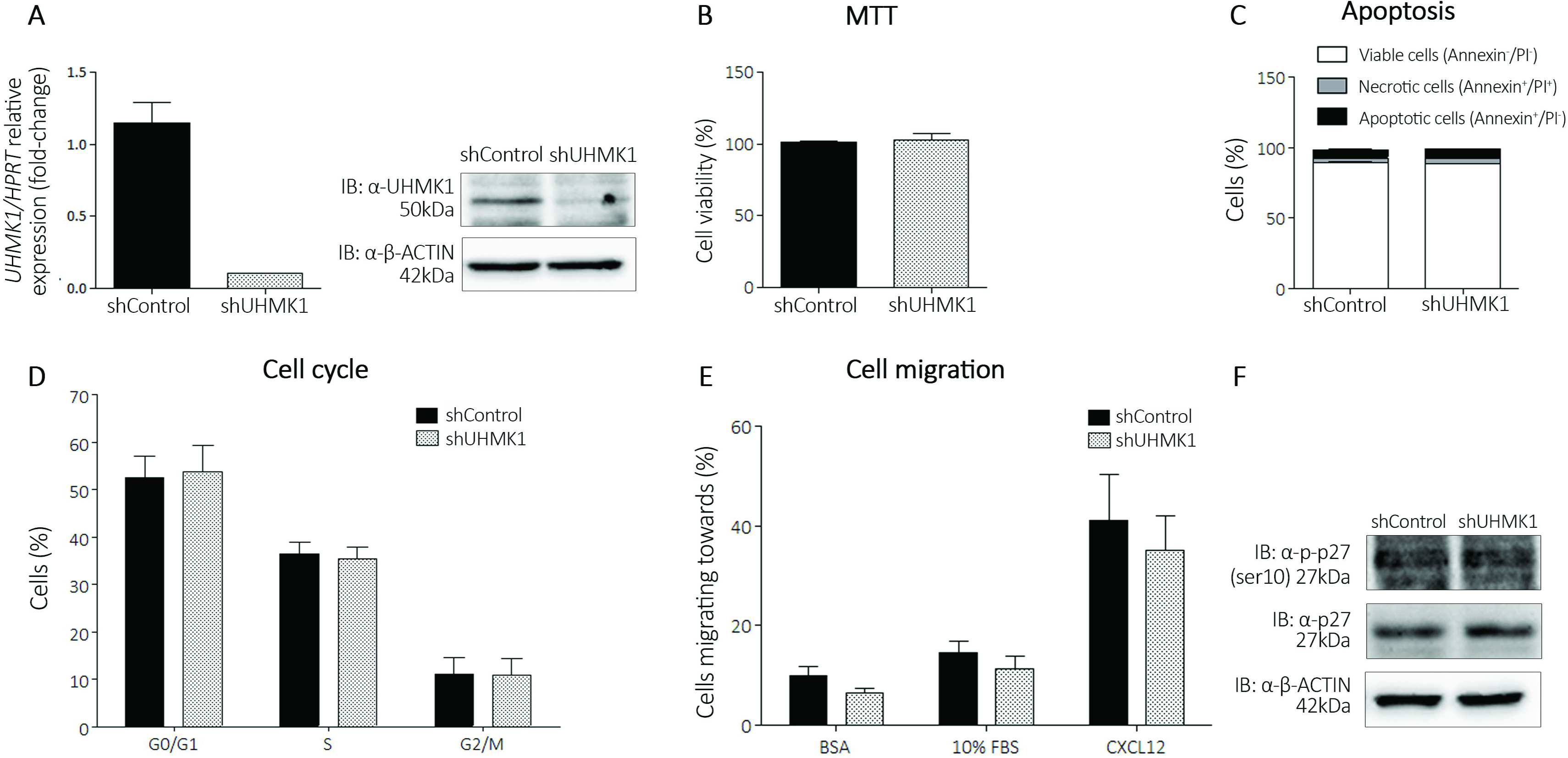
UHMK1 silencing does not affect proliferation and migration of U937 leukemic cells. **(A)** Relative expression of *UHMK1* in control (shControl) and *UHMK1* depleted U937 cells (shUHMK1). *UHMK1* mRNA levels were normalized by *HPRT*. Western blot analysis of shControl and shUHMK1 total cell extracts. Membrane was blotted against anti-UHMK1 and anti-ACTIN, used as loading control. **(B)** Cell viability determined by MTT assay after 48 hours of normal culture conditions. Results are shown as mean ± SD of four independent sixplicate experiments. **(C)** Percentage of viable (annexin V-/PI), apoptotic (annexin V+/PI-) and necrotic cells (annexin V+/PI+), determined by flow cytometry. Results are shown as mean ± SD of three independent duplicate experiments. **(D)** Percentage of cells on different phases of the cell cycle, determined by flow cytometry. Results are shown as mean ± SD of four independent duplicate experiments. **(E)** Migration assay. Number of cells that passed through the pores of a transwell plate towards 0.5% BSA, 10% FBS or CXCL12 containing media after 24 hours exposure to the chemotactic stimuli. The bars represent the number of migrating cells normalized by the input, expressed as percentage. Results are shown as mean ± SD of three independent duplicate experiments. **(F)** p27^KIP^ expression and p27^KIP^(S10) phosphorylation levels in UHMK1 depleted U937 cells. Western blot analysis of shControl and shUHMK1 total cell extracts blotted against anti-p27, anti-phospho-p27(S10) and anti-GAPDH, used as loading control.

UHMK1 silencing did not affect proliferation or viability of U937 cells, as assessed by MTT assay. Also, no difference in apoptosis or the percentage of cells in the different phases of the cell cycle was observed, as assessed by annexin-V/PI staining and analysis of DNA content, respectively (Figure 4B-D). Transwell chemotaxis assay was performed in order to assess the migration of shUHMK1 and shControl cells towards the chemotactic stimulus of FBS (10% Fetal Bovine Serum) and CXCL12. Cell migration was unaffected upon UHMK1 depletion (Figure 4E).

Since UHMK1 is an important regulator of p27^KIP^, we assessed the protein expression and phosphorylation levels of p27^KIP^(S10) on shUHMK1 transduced cells. No difference in the total amount of p27^KIP^ was observed in UHMK1 depleted cells compared to control. Moreover, phosphorylation of p27^KIP^ on S10, a commonly targeted residue by UHMK1, was unaltered upon UHMK1 depletion (Figure 4F). This data confirms that proliferation of the U937 leukemia cells was not affected by silencing UHMK1 protein.

### 3.5. UHMK1 silencing increases clonogenicity of U937 leukemia cells

Colony formation assay was employed to assess the clonogenic potential of the UHMK1 depleted U937 cells on semisolid medium in the absence of growth factors. UHMK1 silenced cells formed 50% more colonies than shControl cells, indicating an augmented autonomous clonal growth of these cells upon UHMK1 depletion (Figure 5).

**Figure 5:**
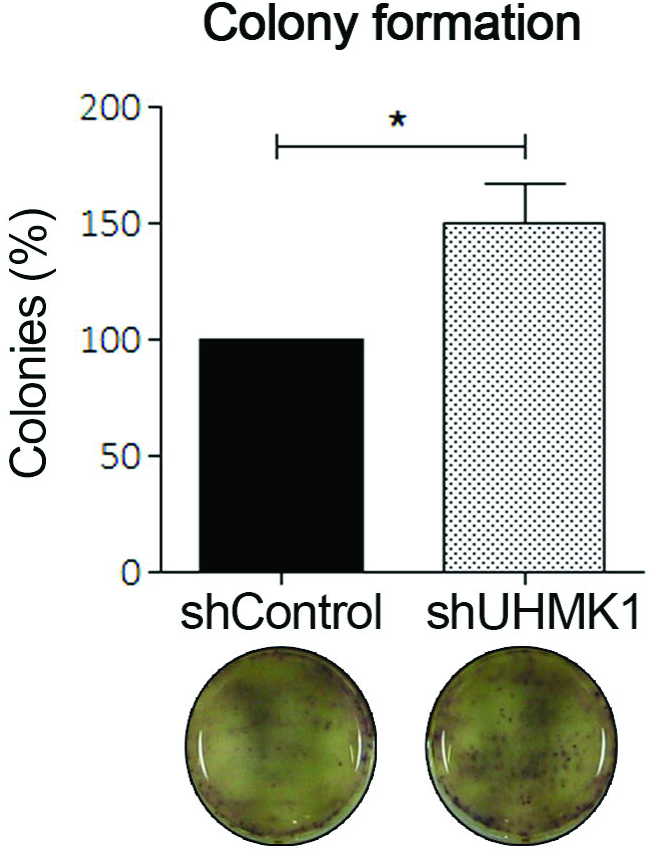
UHMK1 silencing increases clonogenic potential of U937 cells. Percentage of colonies formed after eight days of culture in semi-solid media without growth factors. The bars represent the number of colonies formed normalized by the control and expressed as percentage. Images of representative culturing plates are shown. Results are shown as mean ± SD of six independent triplicate experiments. \**p* < 0.05, Student’s t test.

## 4. Discussion

In order to explore UHMK1 function in the human hematopoietic compartment we performed an extensive analysis of UHMK1 expression in normal and malignant hematopoietic cells, during induced differentiation of primary CD34^+^ cells and leukemia cell lines, and assessed the effect of UHMK1 silencing in the U937 leukemia cells.

Comparing UHMK1 expression in different subpopulations of the bone marrow or peripheral blood, we observed higher levels in the most differentiated cells (lymphocytes CD4^+^, CD8^+^ and CD19^+^) compared to the progenitor enriched subpopulation (CD34^+^) or leukemia cell lines.

Expression analysis on induced differentiation of leukemia cell lines and of primary CD34^+^ cells showed that UHMK1 expression increased upon exposure to a variety of differentiation agents. The KU812 was the only leukemia cell line which did not present UHMK1 upregulation upon differentiation. Of note, KU812 exhibited the highest levels of *UHMK1* transcripts among all leukemia cell lines tested, thus its expression was already elevated prior to erythroid differentiation, which might explain why no increase in the expression was observed. In fact, we show a clear upregulation of UHMK1 when primary human CD34^+^ cells were induced to differentiate into erythrocytes, showing that UHMK1 expression is upregulated in all types of differentiation tested, namely erythroid, megakaryocytic, monocytic and granulocytic differentiation. These results suggest a role of UHMK1 in hematopoietic cell differentiation. A similar suggestion of a possible role for UHMK1 in differentiation has been previously proposed for neuronal cells [9], and more recently for osteoblasts and osteoclasts in bone metabolism [36].

The function of UHMK1 as a regulator of cell cycle progression is well documented. UHMK1 regulates the cyclin dependent kinase inhibitor p27^KIP^ and consequently the cell cycle progression [3]. Thus, it is expected that abnormally elevated UHMK1 activity could be involved in some aspects of tumor development. For this reason, we aimed to investigate *UHMK1* expression on primary cells of patient samples with hematological malignancies. No aberrant expression was observed in patient samples of MDS, AML and ALL compared to healthy donors. However, high expression of *UHMK1* was an independent prognostic factor for increased overall survival in our cohort of patients with MDS. To our knowledge this is the first time that the impact of *UHMK1* expression on MDS survival outcomes has been addressed. Interestingly, UHMK1 is a kinase that phosphorylates the splicing factors SF3b1 and SF1 [4, 5], commonly found mutated in MDS patients [29]. Nevertheless, mutations within *UHMK1* have not been described yet. It is well known that splicing factor function is regulated through reversible phosphorylation events that contribute to assembly of regulatory proteins on pre-mRNA and therefore contributes to the splicing code [37]. It is tempting to speculate that the higher expression of *UHMK1* would impact the regulation and/or function of its substrates SF3B1 and SF1 in MDS patients with better outcome.

Using a lentivirus-mediated shRNA knockdown, we demonstrated that UHMK1 silencing did not affect proliferation or cell cycle progression of U937 leukemia cells. This was supported by the unaltered levels of p27^KIP^ expression and S10 phosphorylation in UHMK1 depleted cells. Similarly, *UHMK1* silencing in breast cancer cells also did not exhibit cell cycle arrest but did when in association with erlotinib [24].

These results were at first confusing since UHMK1 knockdown in cancer cells was expected to reduce p27^KIP^ S10 phosphorylation, stabilize p27^KIP^ and enhanced growth arrest, as observed in other cell line models [3, 20, 23, 38]. On the other hand, cell proliferation and differentiation are two inversely correlated processes and the high levels of UHMK1 expression in differentiated hematopoietic cells seems contradictory to the proliferation promoting function of UHMK1.

It is important to consider that UHKM1 function to overcome growth arrest induced by p27^KIP^ is dependent on mitogenic activation [3]. The fact that UHMK1 depletion did not affect the proliferation of leukemic cells could be explained by the fact that these cells are transformed proliferating, and not quiescent cells arrested in a p27^KIP^ dependent manner. In order to confirm the ability of UHMK1 to overcome p27^KIP^ dependent arrest, we analyzed UHMK1 expression after quiescent PBLs were induced to proliferate with PHA. Western blot analysis revealed a clear upregulation of UHMK1 in proliferating PBLs, compared to quiescent cells. Accordingly, we show massive increase of p27^KIP^ S10 phosphorylation and clear reduction of the total amount of the protein (Supplementary Figure 1). Thus, UHMK1 can induce cell cycle progression in hematopoietic cells in order to release them from cell cycle arrest.

It was shown in an experimental mouse model of vascular wound repair that VMSC Uhmk1^-/-^ cells exhibit increased migratory activity [23]. We did not observe altered migration of UHMK1 silenced leukemia cells assessed by transwell system. The function of UHMK1 in VMSC migration is dependent on its ability to promote STATHMIN phosphorylation and degradation. In that report, the authors did not observe deregulation of STATHMIN in Uhmk1^-/-^ T cells suggesting that STATHMIN based function of Uhmk1 is cell type dependent [23].

Most importantly, UHMK1 silencing boosted colony formation of the U937 cells plated on semisolid culture medium deprived of growth factor. These results suggest that UHMK1 plays a role in the autonomous clonal growth of U937 cells. It is tempting to speculate that the low levels of UHMK1 would render the cells with a more undifferentiated phenotype, and as a consequence turn these cells more clonogenic. Nevertheless, this hypothesis has to be further investigated.

In summary, we show an increased UHMK1 expression in normal and malignant hematopoietic cells induced to differentiate, an increased clonogenicity of UHMK1 depleted leukemia cells and that increased levels of *UHMK1* expression positively impacted prognosis in MDS. Thus, our data suggest that UHMK1 might play a role in hematopoietic cell differentiation.

## Acknowledgements

We thank Roy Edward Larson for English review and Fernanda Soares Niemann, Karla Priscila Ferro and Adriana S. S. Duarte for technical assistance. This research was supported by the Coordenação de Aperfeiçoamento Pessoal de nível Superior (CAPES to IB) and Fundação de Amparo à Pesquisa de São Paulo (FAPESP; 2014/01458-3 to LFA and 2011/51959-0 to STOS). The Hematology and Transfusion Medicine Center-UNICAMP is part of the National Blood Institute (INCT de Sangue CNPq/MCT).

